# Separating distinct structures of multiple macromolecular assemblies from cryo-EM projections

**DOI:** 10.1101/611566

**Authors:** Eric J. Verbeke, Yi Zhou, Andrew P. Horton, Anna L. Mallam, David W. Taylor, Edward M. Marcotte

**Affiliations:** Department of Molecular Biosciences, University of Texas at Austin, Austin, TX 78712, USA; Center for Systems and Synthetic Biology, University of Texas at Austin, Austin, TX 78712, USA; Institute for Cellular and Molecular Biology, University of Texas at Austin, Austin, TX 78712, USA; LIVESTRONG Cancer Institutes, Dell Medical School, Austin, TX 78712, USA

## Abstract

Cryo-electron microscopy is traditionally applied to samples purified to near homogeneity as current reconstruction algorithms are unable to handle heterogeneous mixtures of structures from many macromolecular complexes. We extend on long established methods and demonstrate that relating two-dimensional projection images by their common lines in a graphical framework is sufficient for partitioning distinct protein and multiprotein complexes within the same data set. Using this approach, we first group a large set of synthetic reprojections from 35 unique macromolecular structures ranging from ∼30 – 3000 kDa into individual homogenous classes. We then apply our algorithm on cryo-EM data collected from a mixture of five protein complexes and use existing reconstruction methods to solve multiple three-dimensional structures *ab initio*. Incorporating methods to sort cryo-EM data from heterogeneous mixtures will alleviate the need for stringent purification and pave the way toward investigation of samples containing many unique structures.

## Introduction

Cryo-electron microscopy (cryo-EM) has undergone a revolutionary shift in the past few years. Increased signal in electron micrographs, as a result of direct electron detectors, has allowed for the near-atomic and atomic resolution structure determination of many macromolecules of various shapes and sizes (Kühlbrandt, 2014). These new detectors combined with automated data collection software and improvements in image processing suggest that cryo-EM could be utilized as a high-throughput approach to structural biology. One major obstacle remains: sorting through the immense heterogeneity present in a mixture of tens to hundreds to thousands of macromolecular assemblies.

We and others have shown that cellular extract can be mined for identification of multiple structures (Kastritis et al., 2017; Verbeke et al., 2018). More recently, we showed that it was possible to reconstruct macromolecular machines from the lysate of a single *C. elegans* embryo (Yi et al., 2018). These studies were limited to the identification of only the most abundant and easily identifiable protein and protein–nucleic acid complexes due to a lack of methods to efficiently categorize which two-dimensional (2D) projection images derive from which three-dimensional (3D) assemblies on the basis of their structural features. While a number of 3D classification schemes exist, all failed to produce reliable reconstructions for the majority of particles in these complicated mixtures. This obstacle emphasizes the long-standing need to sort mixtures of structures in addition to their conformational and compositional heterogeneity.

Several methods have been successfully implemented for sorting limited heterogeneity in cryo-EM data. These approaches generally fall into three categories. Currently, the most popular approach for sorting heterogeneity in cryo-EM data utilizes a maximum likelihood estimation to optimize the correct classification of particles into multiple structures (Scheres, 2012; Sigworth, 1998; Sigworth et al., 2010). Another approach is to estimate the covariance in cryo-EM data to search for regions of variability between the models and the data (Katsevich et al., 2015; Liao et al., 2015; Penczek et al., 2006). The last approach involves computing similarities between projection images in the data before applying clustering methods to separate the data into homogenous subsets (Aizenbud and Shkolnisky, 2016; Herman and Kalinowski, 2008; Shatsky et al., 2010). All of these approaches have been demonstrated on samples containing a primary structure with multiple conformations or variable subunits. However, little work has been done for sorting heterogeneous samples containing multiple distinct structures.

Here, we develop a pipeline for building 3D reconstructions from a mixture of distinct particles by first grouping 2D projections into discrete, particle-specific classes using the principles of common lines and a novel graphical clustering framework. We demonstrate our method by partitioning reprojections from 35 previously solved X-ray crystal structures into their correct groups. Furthermore, we applied this pipeline to an experimental set of cryo-EM micrographs containing a mixture of several macromolecular complexes. We were able to reconstruct multiple 3D structures after our clustering, improving on classification of all particles simultaneously using current 3D reconstruction software. These results are a necessary first step for moving cryo-EM towards high-throughput structural biology.

## Results

### Classifying projection images from multiple structures

A major challenge facing “shotgun”-style cryo-EM is to reconstruct models from projection images arising from multiple distinct structures present in a mixture. To overcome this obstacle, we sought a method to computationally group heterogeneous projection images into discrete classes that each derive from the same structure. Two-dimensional projection images from the same asymmetric object can be related to each other if there is prior information of the three-dimensional object (i.e. an initial model) using projection-matching algorithms. One approach that circumvents the need for a starting model is to relate the 2D projection images based on common lines (Van Heel, 1987), derived from the projection-slice theorem, which states that any two 2D projections of the same 3D object must share a 1D line projection in common. In order to partition projection images into homogenous subsets, we developed an algorithm for detecting **<u>S</u>**hared **<u>L</u>**ines **I**n **<u>C</u>**ommon **<u>E</u>**lectron **<u>M</u>**aps (SLICEM). Using this algorithm, we score the similarity of 1D line projections between sets of 2D projection images without knowledge of the underlying 3D objects. Subsequently, these similarity scores can be put into a graphical framework and clustering algorithms can be applied to group related 2D projection images for subsequent 3D reconstructions (Figure 1).

**Figure 1.**
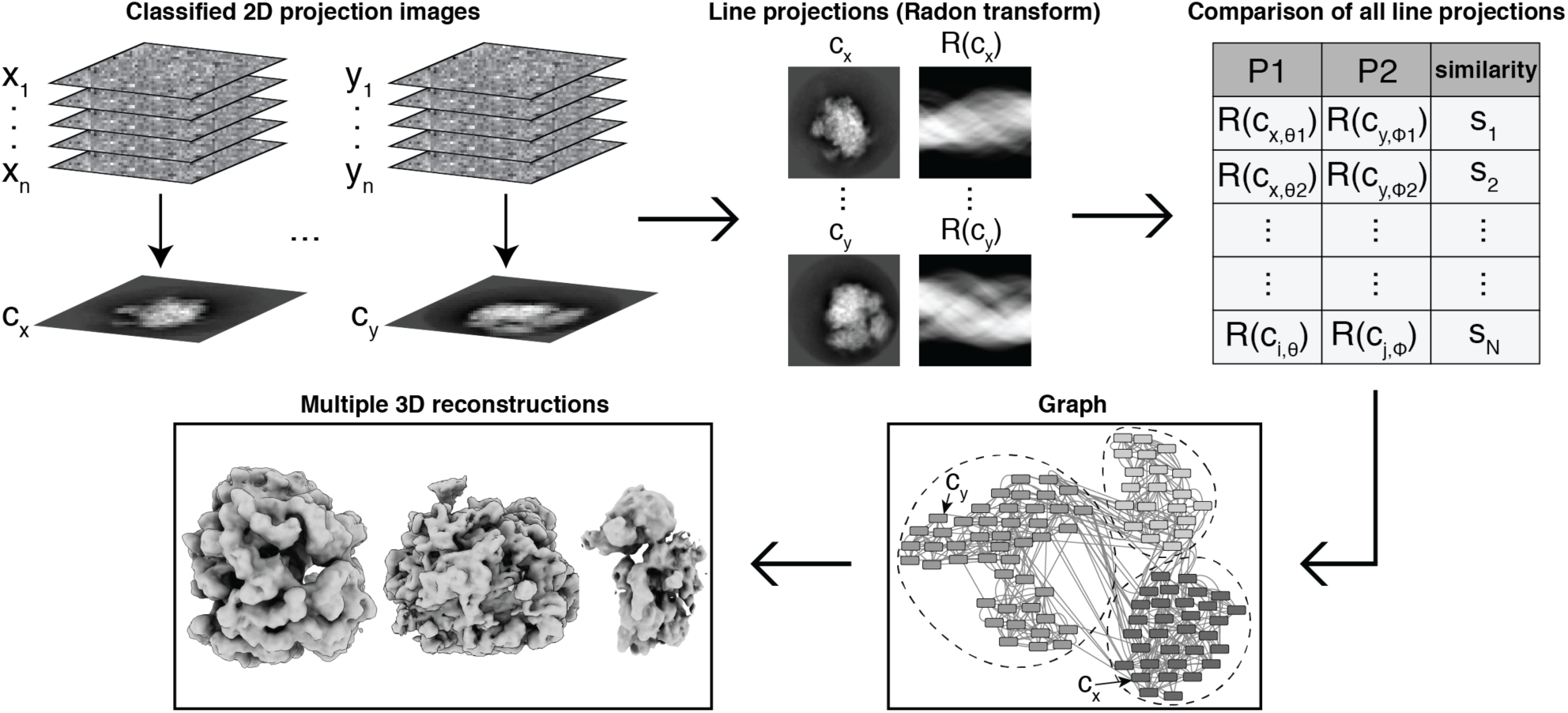
Computational pipeline for SLICEM. Individual particle images are averaged after reference-free 2D alignment and classification. Using a Radon transform, 1D line projections are created from the 2D class averages (referred to as 2D projections). Each 1D line projection from every 2D projection is then scored for similarity. The top scores between each projection are used to form edges connecting 2D projections that have a similar 1D line projection to form a graph. 2D projection images are then partitioned into groups belonging to the same putative structure using a community detection algorithm. Individual particle images belonging to each 2D projection within a community are subjected to *ab initio* 3D reconstruction.

### Synthetic data

To test our approach using SLICEM, we generated synthetic reprojections from 35 previously solved X-ray crystal structures (see Methods) (Figure S1). The structures ranged in molecular weight from ∼30 – 3000 kDa. Each structure was low-pass filtered to 9 Å and uniformly reprojected to create 12 2D projection images (Ludtke et al., 1999). Next, we combined reprojections from all models to simulate ideal 2D class averages from a heterogeneous cryo-EM dataset. The similarity of 1D line projections from each image was scored using 6 different metrics (see Methods). The precision and recall of correctly pairing 2D projection images from the same 3D structures was computed in order to determine the performance of each metric, and cosine similarity was determined to be the top performing metric (Figure 2A).

**Figure 2.**
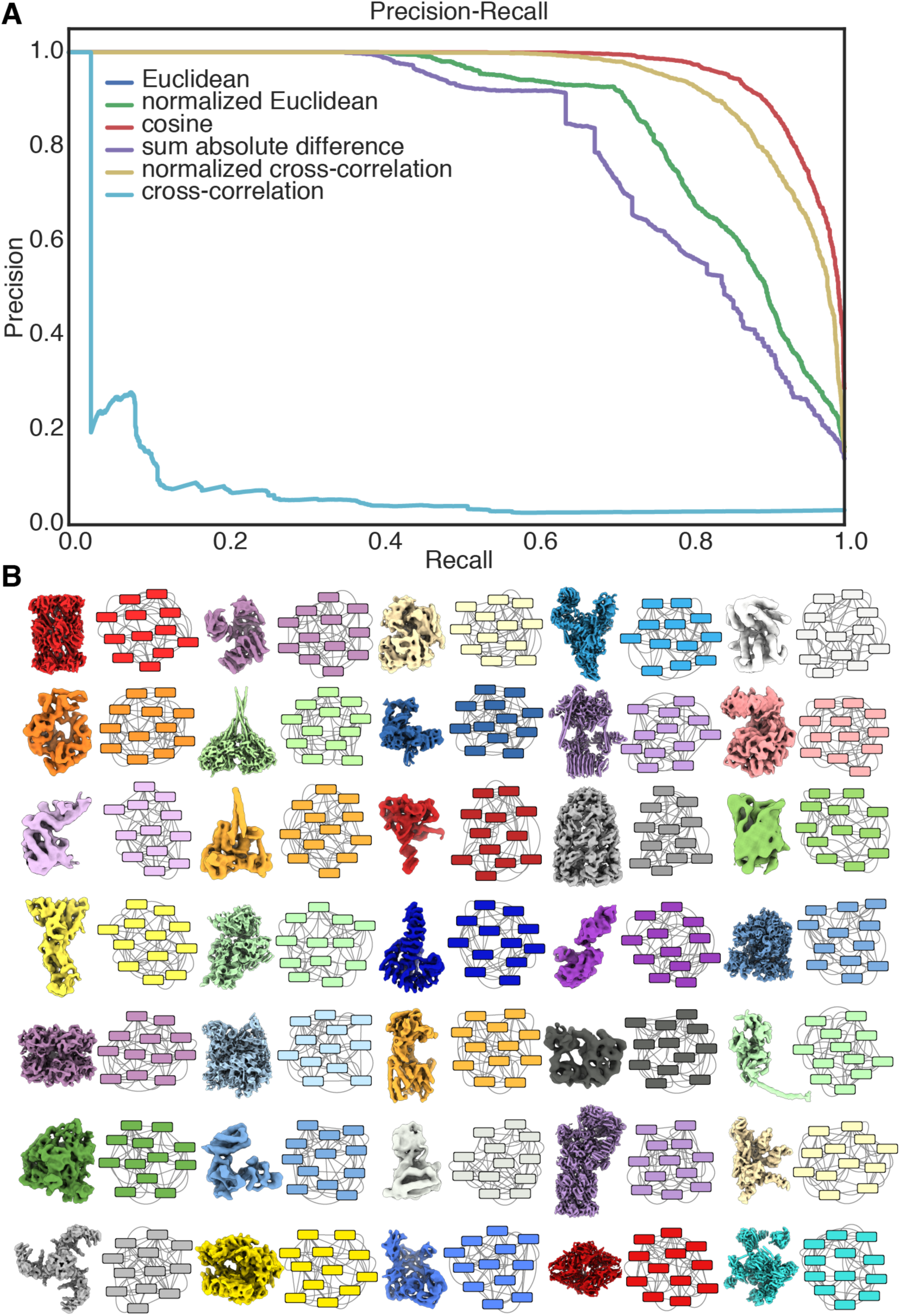
Separating mixtures of synthetic 2D reprojections. Synthetic reprojections were generated from 35 distinct X-ray crystal structures low-pass filtered to 9 Å from complexes ranging in molecular weight from ∼30 – 3000 kDa, prior to separation using SLICEM. (A) Precision-recall plot ranking 6 different metrics at scoring the similarity between 1D line projections from each 2D reprojection. (B) Network output displaying communities of 2D reprojection images determined using SLICEM. Each node represents a 2D reprojection with 5 connecting edges to the most similar reprojections as scored using Euclidean distance. The color of each node matches the structure from which it was reprojected (shown as a surface).

In order to identify sets of 2D projection images from the same 3D particles, we constructed a network from the comparisons between projection images as follows: Each 2D projection image was represented as a node in a directed graph, with each node connected by edges to the nodes corresponding to the 5 most closely-related 2D projection images based on the similarity of their 1D line projections. While the top-scoring metric in our precision/recall analysis was cosine similarity, the network generated from the Euclidean distance similarity most clearly showed communities (clusters of 2D projections) correctly partitioned by 3D structure (Figure 2B). These results show that partitioning 2D projection images by scoring their common lines is a powerful, unsupervised approach for sorting cryo-EM data from distinct 3D structures within a heterogeneous mixture.

### Cryo-EM on a mixture of protein complexes

After validating our SLICEM algorithm on a synthetic dataset, we performed cryo-EM on an experimental mixture of structures and tested our approach as a proof-of-principle. Our experimental mixture consisted of 40S, 60S and 80S ribosomes, apoferritin and β-galactosidase. We collected ∼2,400 images and used a template-based particle picking scheme to select ∼523,000 particles from the entire data set (Roseman, 2004). Raw micrographs showed a mixture of disperse particles with varying size and shape (Figure S2). We then performed 2D classification on the entire set of particles using RELION (Scheres, 2012). After 1 round of filtering junk particles, the remaining ∼203,000 particles were sorted into 100 classes using RELION. The class averages contained many characteristic ribosome projections and had distinct structural features (Figure S2). We were unable to identify any β-galactosidase particles in our collected images.

We then applied our SLICEM algorithm to the 100 2D class averages. The identity of each class average was manually annotated, where it was easily recognizable, to assess whether our algorithm was correctly separating the 2D projection images from our heterogeneous mixture (Figure 3). Based on these manual annotations, we again tested the 6 different metrics in a precision-recall framework to determine which metric performed better on experimental data (Figure S3). Interestingly, the Euclidean distance and sum of the absolute difference scoring metric significantly outperformed the cosine similarity. Using the sum of the absolute difference scoring metric, the network naturally partitioned into 3 distinct communities, one for each ribosome, prior to employing any community detection algorithms (Figure 3). As part of our algorithm, we evaluated two community detection methods, edge betweenness and walktrap, to determine if the network should be further subdivided (Latapy and Pons, 2004; Newman and Girvan, 2004). We chose to use community detection algorithms to prevent biasing the data by choosing the number of output clusters we expected. Using our SLICEM algorithm, we were able to correctly separate 2D projection images from 3 large, asymmetric macromolecular complexes from the same mixture.

**Figure 3.**
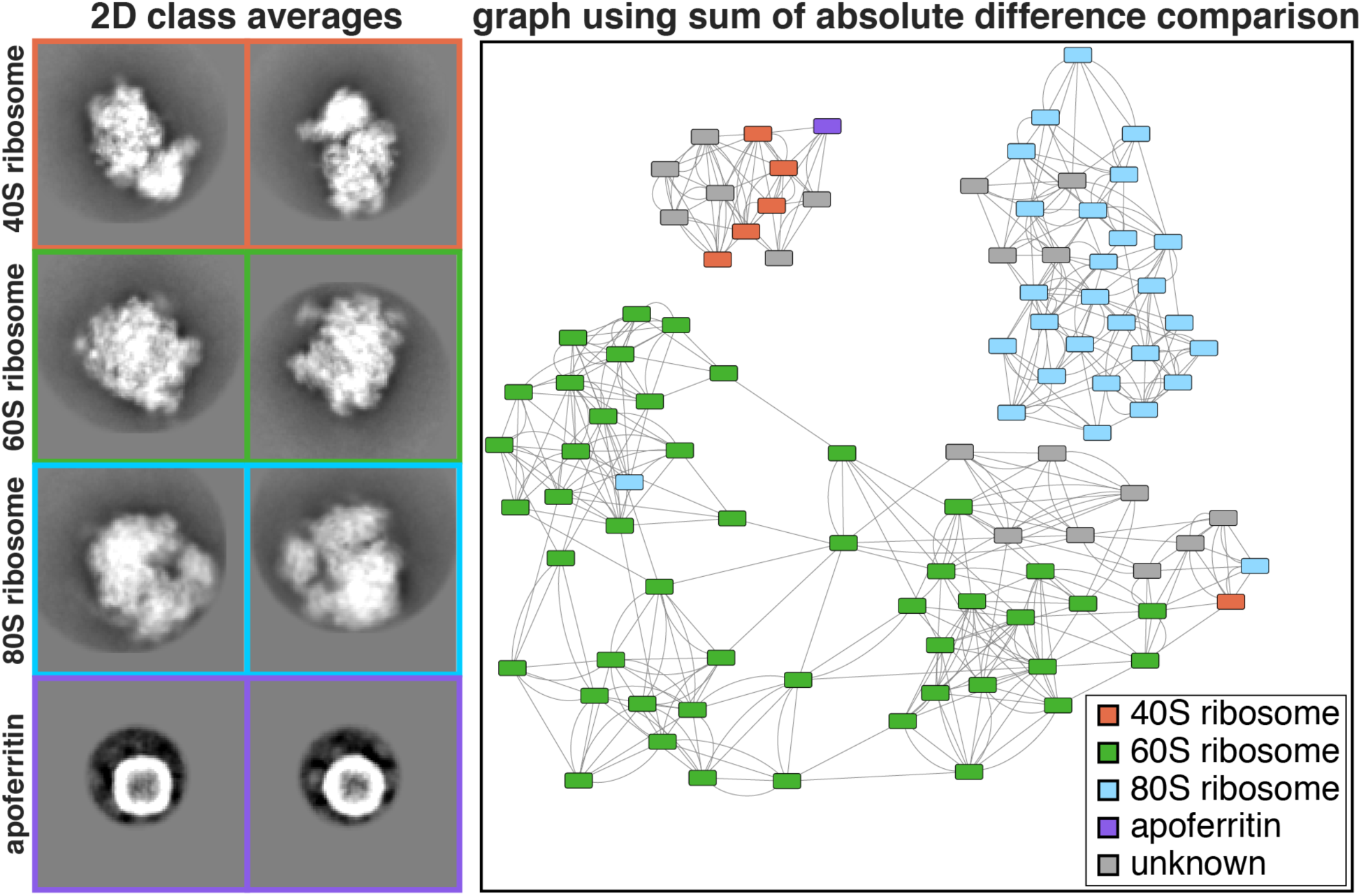
Experimental 2D class averages and resulting network. Cryo-EM data was collected on a mixture of 5 protein and protein-nucleic acid complexes. (A) Representative 2D class averages of the 4 complexes identified in the mixture. The identity of each class average was manually annotated were it could be easily identified. The class average corresponding to apoferritin was further subdivided into multiple classes for visualization. (B) Network generated using SLICEM on the 100 2D class averages scored using the sum of the absolute difference metric. Nodes representing each 2D class averages are colored by their putative structural identity. The width of the box corresponds to 422 Å.

### Relating summed pixel intensity to molecular weight

Apart from partitioning 2D projection images into homogenous subsets for 3D reconstruction, one additional goal was to determine the identity of each projection image. In previous studies, we and others have leveraged mass spectrometry data to help identify electron microscopy reconstructions from a heterogeneous mixture, such as cell lysate, where the architecture of every protein or protein complex is not known (Kastritis et al., 2017; Verbeke et al., 2018). However, this combined MS-EM approach was only useful for identifying highly abundant and easily recognizable structures.

To provide evidence of macromolecular identity from the electron maps, we calculated the sum of pixel intensities in each manually annotated 2D class average as a proxy for molecular weight (Figure 4). We found that each of the three ribosomes and apoferritin had unique summed pixel intensities that could be used to distinguish the class averages. A least-squares fit to the mean of the summed pixel intensities showed a linear relationship between summed pixel intensity and protein molecular weight. The summed pixel intensities were therefore used as an additional filtering step by removing nodes in communities whose summed pixel intensities were outliers in that community. Using this filtering step, the apoferritin class average was removed from the community containing predominantly 40S ribosome reprojections. Our data suggest that, given an appropriate set of standards, summed pixel intensity can be correlated to molecular weight. Thus, summed pixel intensity could be useful in narrowing down the possible identities for a set of electron densities, when combined with sequence information.

**Figure 4.**
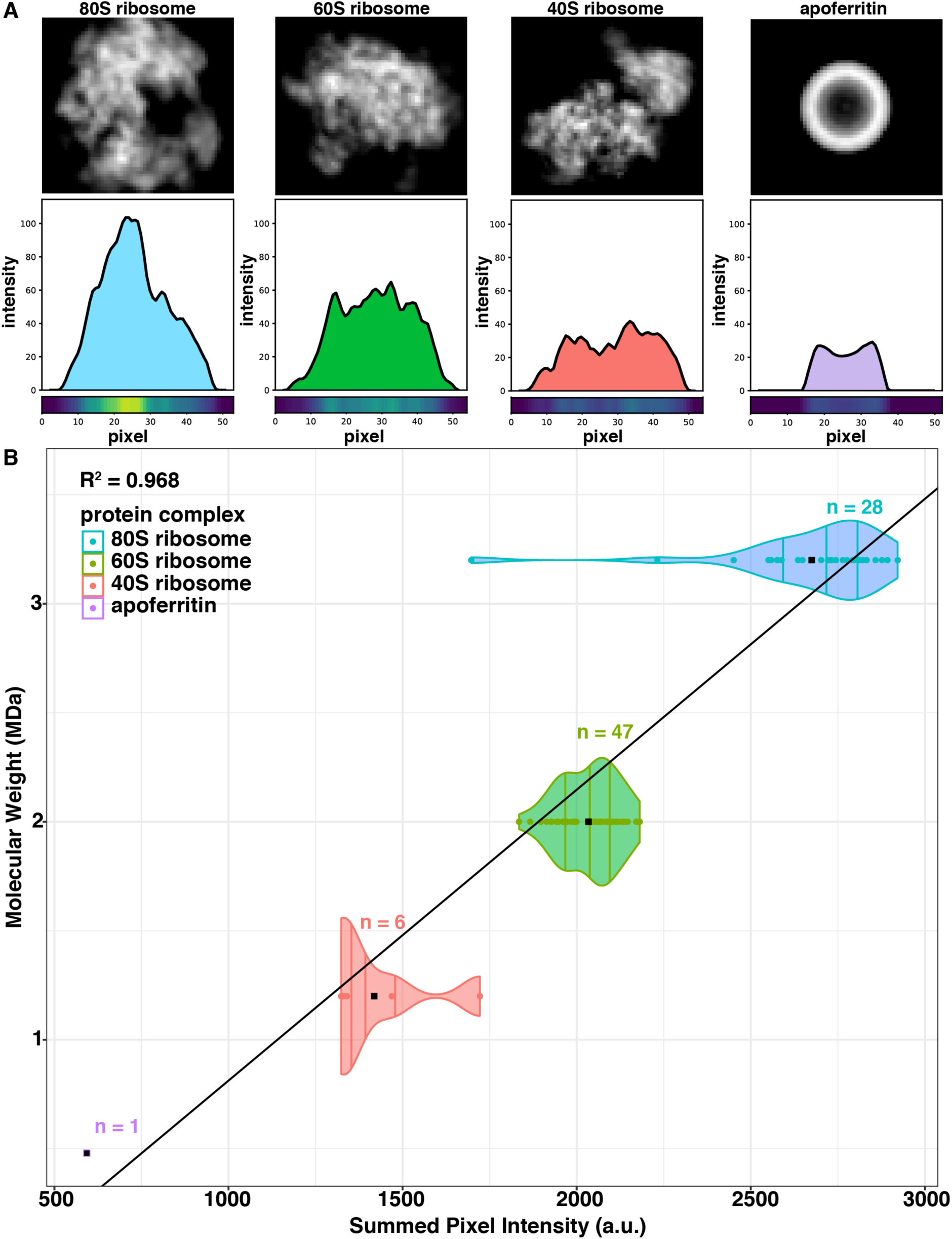
Summed pixel intensities of 2D class averages correlate to molecular weight. (A) 2D to 1D projections for representative 2D class averages of each structure present in the mixture. 1D projection plots show the line profile for a single projection of each 2D class average. Pixel heat maps show the intensity of the line profile at each pixel. (B) Distribution of the summed 1D projection pixel intensities, or integration of the 1D line profiles, calculated for each 2D class average. Summed pixel intensities for each manually identified 2D class average are plotted against their respective molecular weight. Black points are the mean summed pixel intensity for each structure.

### 3D classification of a mixture of protein complexes

The ultimate goal of our pipeline is to reconstruct 3D models from our output of clustered 2D projection images. We chose to use cryoSPARC for 3D reconstructions because it can perform heterogeneous reconstruction without *a priori* information on structure or identity (Punjani et al., 2017). We used the particles from each of our 3 distinct communities in addition to the isolated apoferritin node for *ab initio* reconstruction in cryoSPARC (Figure 5). The cluster containing primarily 40S ribosome particles was split into two classes to filter the additional junk particles present in the community. Comparison of our models reconstructed after clustering to the models produced using the entire data set as input for *ab initio* reconstruction in cryoSPARC with 4 classes (one for each protein complex in the mixture) showed our pre-sorting procedure improved the resulting structures (Figure 5). In particular, we were able to build an apoferritin model that was missed in the 3D classification of all particles from cryoSPARC. Our 80S model also shows a more complete density for the small subunit than its counterpart in the model created without clustering. We also observe that changing the number of classes using *ab initio* reconstruction in cryoSPARC had a substantial impact on the quality of classification (Figure S4).

**Figure 5.**
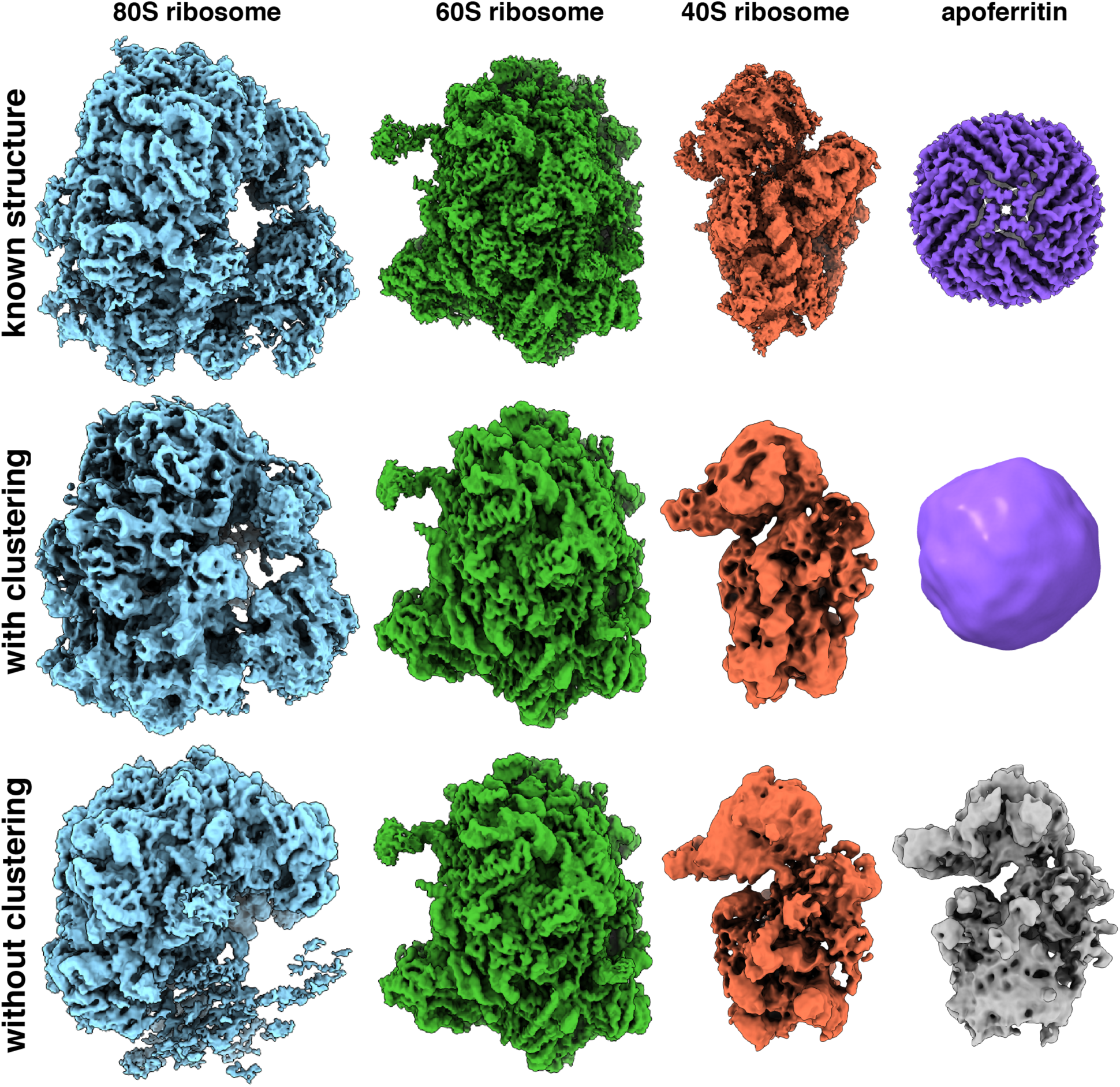
*Ab initio* structures from an experimental mixture. (A) High-resolution structures of the 80S ribosome EMD-2858 (Cianfrocco and Leschziner, 2015), 60S ribosome EMD-2811 (Shen et al., 2015), 40S ribosome EMD-4214 (Scaiola et al., 2018) and apoferritin EMD-2788 (Russo and Passmore, 2014). (B) 3D models of the 80S ribosome, 60S ribosome, 40S ribosome and apoferritin generated by sorting particles using SLICEM prior to *ab initio* 3D reconstruction in cryoSPARC. (C) 3D models generated using *ab initio* reconstruction to generate 4 classes in cryoSPARC without pre-sorting particles using SLICEM.

Each model was refined and evaluated using the gold-standard 0.143 Fourier shell correlation criterion (Figure S5). We obtained easily identifiable 40S, 60S, and 80S ribosome structures at 12, 4, and 5.4 Å resolution, respectively. We were also able to reconstruct the smaller, more compact apoferritin at 19 Å resolution. Notably, the 40S and 80S models contain streaks in one dimension, indicating that we are missing several orientations of the particles. We attribute this to preferential orientation of the particles in ice, rather than an inability of our algorithm to properly sort particles into correct communities. Together, these results demonstrate a functioning pipeline for sorting 2D projection images from a heterogeneous mixture of 3D structures, allowing for single particle EM to be applied to samples containing multiple proteins or protein complexes. Importantly, aside from choosing the most appropriate similarity measure, our approach is fully unsupervised, requiring no user defined estimate of the number of existing classes.

## Discussion

As cryo-EM continues to rapidly advance, one potential application would be to perform high-throughput structural biology. The ability to sort and classify heterogeneous mixtures will become a necessary feature. One advantage of this approach would be to study closer-to-native proteins directly from cell lysate without the need to purify or alter the sample. Currently, handling compositional and conformational heterogeneity is a major challenge for the EM field, usually requiring expert, time-consuming steps. In this study, we present an unsupervised algorithm, SLICEM, which extends on previous methods and demonstrates that sorting 2D projection images based on the similarity of their common lines is capable of correctly clustering 2D projection images from a mixture of protein and protein-nucleic acid complexes. We first demonstrate that the algorithm successfully sorts a synthetic dataset of reprojections created from 35 unique macromolecular structures. Next, we show the same algorithm can successfully partition 2D projection images from an experimental data set containing multiple macromolecular complexes. Pre-sorting 2D projection images prior to 3D classification allows current reconstruction algorithms to be employed on datasets that would otherwise be too complex.

Although we demonstrated the feasibility of our approach on synthetic and experimental data, we acknowledge that there are several limitations. In particular, our algorithm relies on the quality of upstream 2D alignment, classification and averaging. As we observed during 2D classification of our cryo-EM data, all apoferritin particles were grouped into a single class average. However, during our network generation step, each class average is given multiple edges to the most similar classes, forcing the single apoferritin class average to have multiple spurious edges. This error will occur any time the number of class averages of a given structure is less than the number of edges used in the graph. Future modifications to the algorithm could include searching for symmetric class averages, where this error is more likely to occur, and removing them prior to community detection.

As we move cryo-EM towards structural determination of heterogeneous mixtures, several other technical challenges will emerge, such as universal freezing conditions. In our mixture of 5 macromolecular complexes, we were unable to easily find freezing conditions that accommodated all proteins. The result was a mixture missing β-galactosidase and containing orientation preferences for the 40S and 80S ribosome. However, previous work has produced e.g. high-resolution structures of fatty-acid synthase from fractionated cell lysate, suggesting it is possible to find suitable cryo-conditions for solutions containing many macromolecular species (Kastritis et al., 2017). An additional challenge will be developing particle picking algorithms specifically for mixtures, where the particle shape may be unknown and, perhaps more importantly, non-uniform. While in this study we used a template picking scheme, future studies with mixtures of unknown composition will require more sophisticated approaches.

An expert might be able to manually sort the class averages from our cryo-EM data set; however, as mixtures grow in complexity, manual sorting will certainly become infeasible. Introducing algorithms such as SLICEM will provide an unbiased way to group 2D projection images and can be easily implemented in conjunction with a variety of image processing and 3D reconstruction packages. One additional utility of this algorithm could be to remove junk class averages from data in a semi-supervised manner by removal of communities of projection images that do not appear to have structural features. Our approach for sorting mixtures of structures combined with previous approaches for sorting conformational heterogeneity could be a powerful tool for deep classification. Development of methods to sort mixtures of structures in single particle cryo-EM will allow us to solve more structures in parallel and alleviate time-consuming protein purification and sample preparation.

## Materials and Methods

### Synthetic data generation

The following list of PDB entries were used to create the dataset of synthetic reprojections (1A0I, 1HHO, 1NW9, 1WA5, 3JCK, 5A63, 1A36, 1HNW, 1PJR, 2FFL, 3JCR, 5GJQ, 1AON, 1I6H, 1RYP, 2MYS, 3VKH, 5VOX, 1FA0, 1JLB, 1S5L, 2NN6, 4F3T, 6B3R, 1FPY, 1MUH, 1SXJ, 2SRC, 4V6C, 6D6V, 1GFL, 1NJI, 1TAU, 3JB9, 5A1A). Each PDB entry was low-pass filtered to 9 Å and converted to a 3D EM density using ‘pdb2mrc’ in EMAN (Ludtke et al., 1999). These densities were then uniformly reprojected using ‘project3d’ in EMAN to create 12 2D reprojections for each structure (Ludtke et al., 1999). Reprojections were centered in 350 Å boxes.

### Purification of apoferritin and β-galactosidase

Size-exclusion chromatography was performed at 4 °C on an AKTA FPLC (GE Healthcare). Approximately 10 mg of apoferritin (Sigma A3660-1VL) and 5 mg of β-galactosidase G5635-5KU were independently applied to a Superdex 200 10/300 GL analytical gel filtration column (GE Healthcare) equilibrated in 20 mM HEPES KOH, 100 mM potassium acetate, 2.5 mM magnesium acetate, pH 7.5 at a flow rate of 0.5 mL min-1. Fractions were collected every 0.5 mL.

### SLICEM Algorithm

Our algorithm consists of five main steps: (1) Extracting 2D class average signal from background, (2) Generating 1D line projections from the extracted 2D projection images, (3) Scoring the similarity of all pairs of 1D line projections, (4) Building a nearest-neighbors graph of the 2D class averages and (5) Partitioning communities within the graph.

#### (1) Extracting 2D class averages from background

The input to our algorithm is a set of centered and normalized 2D class averages. We then extract the centered region of positive pixels values from the zero-mean normalized images to remove background signal and extra densities that might be present in a class average.

#### (2) Generating 1D line projections from extracted 2D projection images

Each extracted class average is projected into 1D over 360 degrees in 5 degree intervals by summing the pixel values along the projection axis. The 1D line projections are then independently zero-mean normalized if the normalized cross-correlation or normalized Euclidean scoring metric are selected.

#### (3) Scoring the similarity of all pairs of 1D line projections

To score the similarity of the 1D line projections we considered 6 different scoring metrics: Euclidean distance, normalized Euclidean distance, cosine similarity, sum of the absolute difference, cross-correlation and normalized cross-correlation. For the non-cross-correlation metrics, the similarity of the 1D line projections is calculated for translations of the smaller 1D projection across the larger 1D projection if there is a difference in projection size, analogous to the ‘sliding’ feature of cross-correlations. The optimum score during the translations is kept for each pair of 1D projections. After pairwise scoring of all 1D line projections, the similarity between each pair of 2D class averages is defined by their respective highest scoring 1D line projections.

#### (4) Building a nearest-neighbors graph of the 2D class averages

SLICEM then constructs a directed graph using the similarity scores calculated for each pair of 2D class averages. Each node (2D class average) is connected to the 5 most similar (top scoring) 2D class averages. Each edge is assigned a weight computed as a z-score relative to all scores for a given 2D class average.

#### (5) Partitioning communities within the graph

The resulting graph is then subdivided using a community detection algorithm. Specifically, we evaluated the edge-betweenness and walktrap algorithms to define clusters in the graph. Then, the median absolute deviation of summed pixel intensities for each node is calculated to remove outliers from the cluster. The final set of nodes in a cluster is then used as input for 3D reconstruction in cryoSPARC.

#### Cryo-EM grid preparation and data collection

C-flat holey carbon grids (CF-1.2/1.3, Protochips Inc.) were pre-coated with a thin layer of freshly prepared carbon film and glow-discharged for 30 seconds using a Gatan Solarus plasma cleaner before addition of sample. 2.5 μl of a mixture of 75 nM 40S ribosome, 150 nM 60S ribosome, 50 nM 80S ribosome, 125 nM apoferritin and 125 nM β-galactosidase were placed onto grids, blotted for 3 seconds with a blotting force of 5 and rapidly plunged into liquid ethane using a FEI Vitrobot MarkIV operated at 4 °C and 100% humidity. Data were acquired using an FEI Titan Krios transmission electron microscope (Sauer Structural Biology Laboratory, University of Texas at Austin) operating at 300 keV at a nominal magnification of ×22,500 (1.1 Å pixel size) with defocus ranging from −2.0 to −3.5 μm. The data were collected using a total exposure of 6 s fractionated into 20 frames (300 ms per frame) with a dose rate of ∼8 electrons per pixel per second and a total exposure dose of ∼40 e^−^ Å^−2^. A total of 2,423 micrographs were automatically recorded on a Gatan K2 Summit direct electron detector operated in counting mode using the MSI Template application within the automated macromolecular microscopy software LEGINON (Suloway et al., 2005).

### Cryo-EM data processing

All image pre-processing was performed in Appion (Lander et al., 2009). Individual movie frames were aligned and averaged using ‘MotionCor2’ drift-correction software (Zheng et al., 2017). These drift-corrected micrographs were binned by 8, and bad micrographs and/or regions of micrographs were removed using the ‘manual masking’ command within Appion. A total of 522,653 particles were picked with a template-based particle picker using a reference-free 2D class average from a small subset of manually picked particles as templates. The contrast transfer function (CTF) of each micrograph was estimated using CTFFIND4 (Rohou and Grigorieff, 2015). Selected particles were extracted from micrographs using particle extraction within RELION (Scheres, 2012) and the EMAN2 coordinates exported from Appion. Two rounds of reference free 2D classification with 100 classes for each sample were performed in RELION to remove junk particles, resulting in a clean stack of 202,611 particle images.

## Author Contributions

E.J.V. developed the code and performed all experiments. Y.Z. prepared samples and collected the cryo-EM data. A.P.H. helped refine the code. A.L.M. helped with protein purification. E.J.V., D.W.T, and E.M.M. conceived of the experiments, analyzed the data, and wrote the manuscript. D.W.T. and E.M.M. supervised and obtained funding for the work. All authors commented on the manuscript.

## Acknowledgements

We thank S. Musalgaonkar and A. Johnson for providing 40S and 60S yeast ribosomes; A. Dai for expert cryo-EM assistance at the Sauer Structural Biology Laboratory at UT Austin; and members of the Taylor and Marcotte laboratories for helpful discussions. D.W.T. is a CPRIT Scholar supported by the Cancer Prevention and Research Institute of Texas (RR160088) and an Army Young Investigator supported by the Army Research Office (WW911NF-19-10021). This work was supported in part by Welch Foundation Research Grants F-1938 (to D.W.T.) and F-1515 (to E.M.M.), Army Research Office Grant (W911NF-15-0120) (to D.W.T.), Robert J. Kleberg, Jr. and Helen C. Kleberg Foundation Medical Research Award (to D.W.T.) and grants from the National Institutes of Health (GM122480, DK110520, HD085901) to E.M.M..

## Data Availability

The cryo-EM reconstructions of the 40S, 60S, 80S, and apoferritin have been deposited in the Electron Microscopy Databank with accession codes EMD-20109, EMD-20110, EMD-20111 and EMD-20112, respectively. The motion-corrected sum micrographs have been deposited into EMPIAR with accession code EMPIAR-10268. Computer code for SLICEM is available at https://github.com/marcottelab/SLICEM.

## Supplemental Information

**Figure S1.**
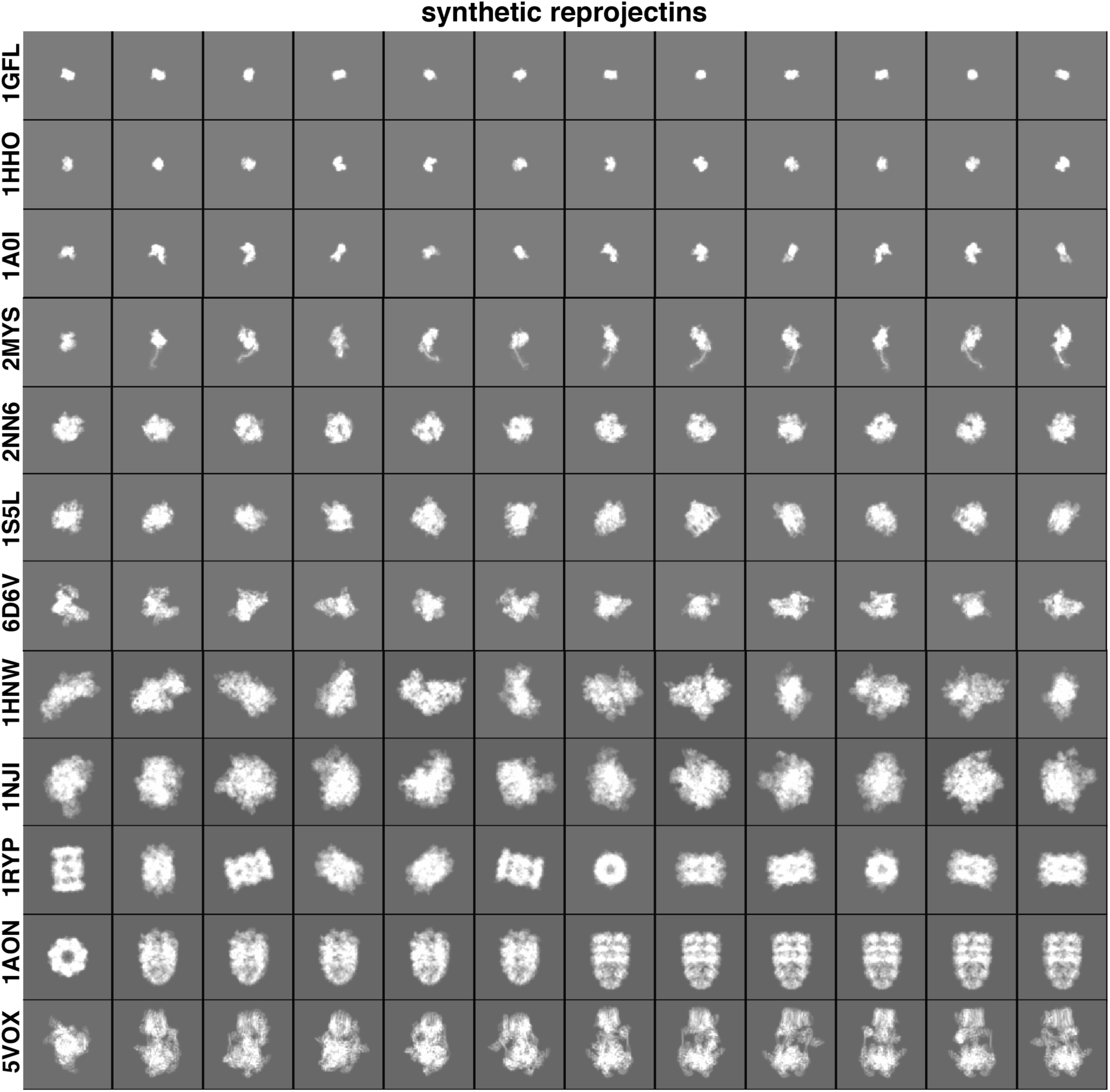
2D reprojections from synthetic dataset. Subset of 2D reprojections from 12 of the 35 structures in our synthetic dataset. Box size corresponds to 300 Å.

**Figure S2.**
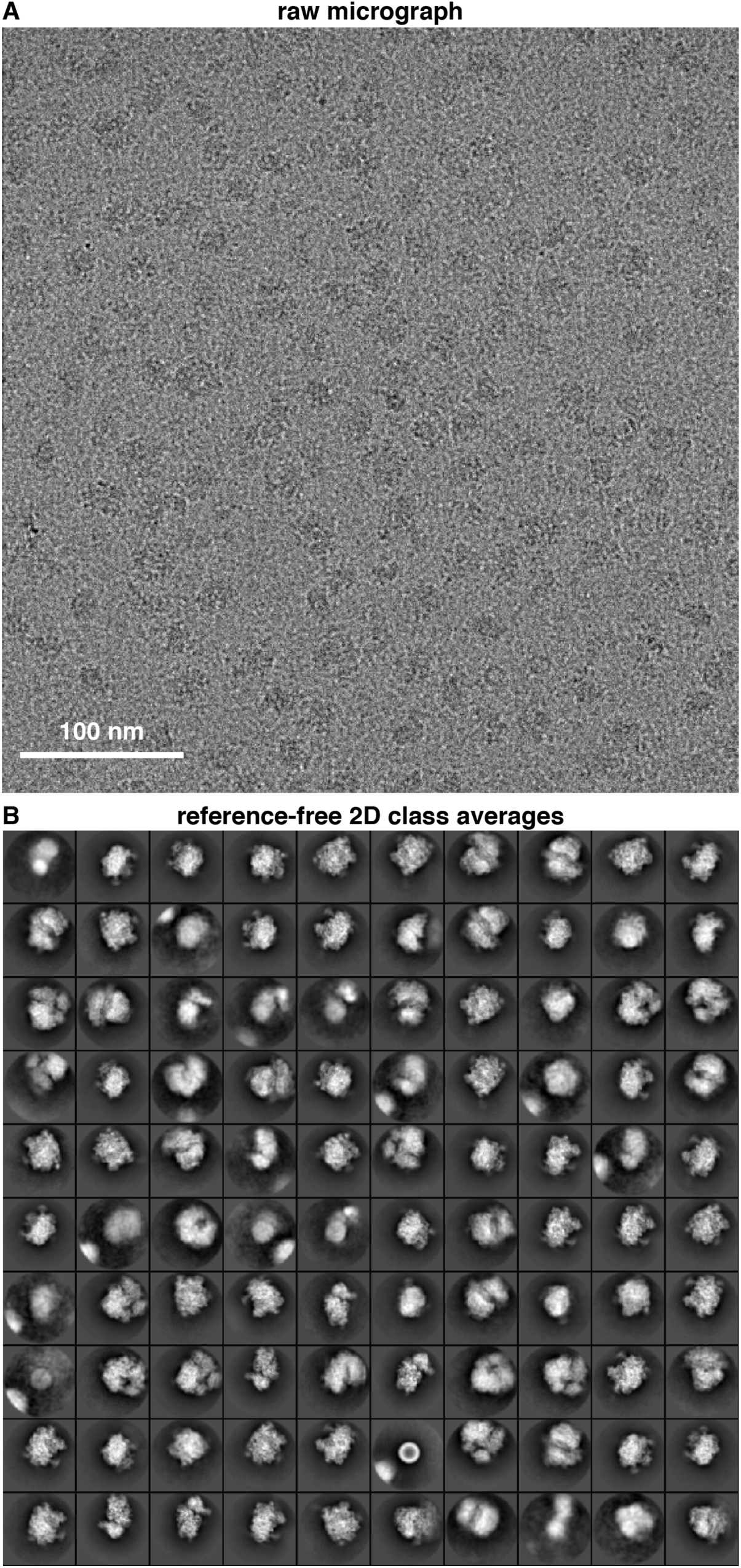
2D classification of particles using RELION. (A) Representative raw micrograph of a mixture containing 40S, 60S and 80S ribosomes, apoferritin and β-galactosidase. (B) Reference-free 2D class averages generated using RELION of ∼203,000 template-picked particle images. Box size corresponds to 422 Å.

**Figure S3.**
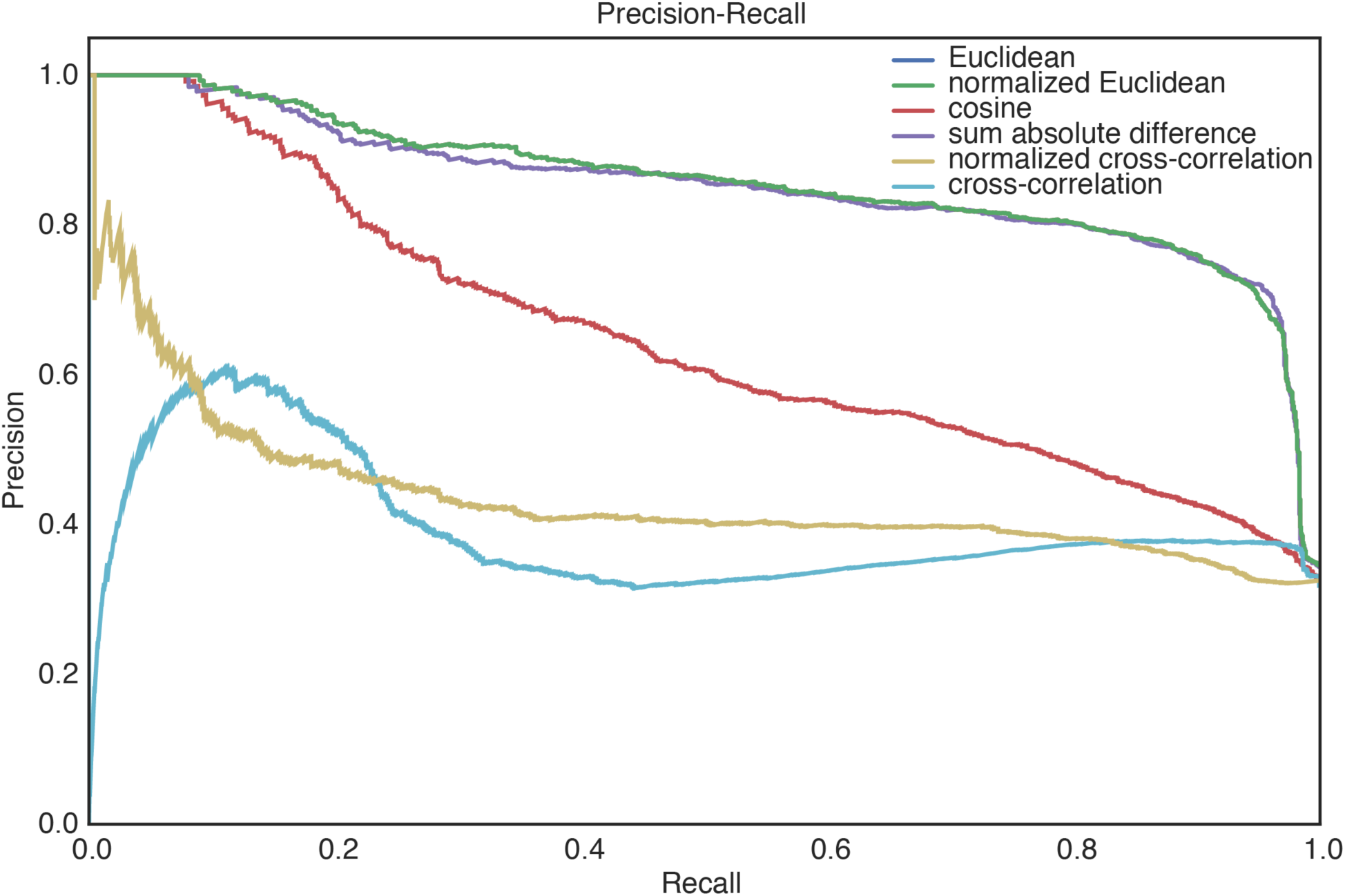
Precision-recall curves for experimental cryo-EM data. Precision-recall plot displaying 6 different metrics for scoring the similarity between 1D line projections from the entire set of 2D class averages.

**Figure S4.**
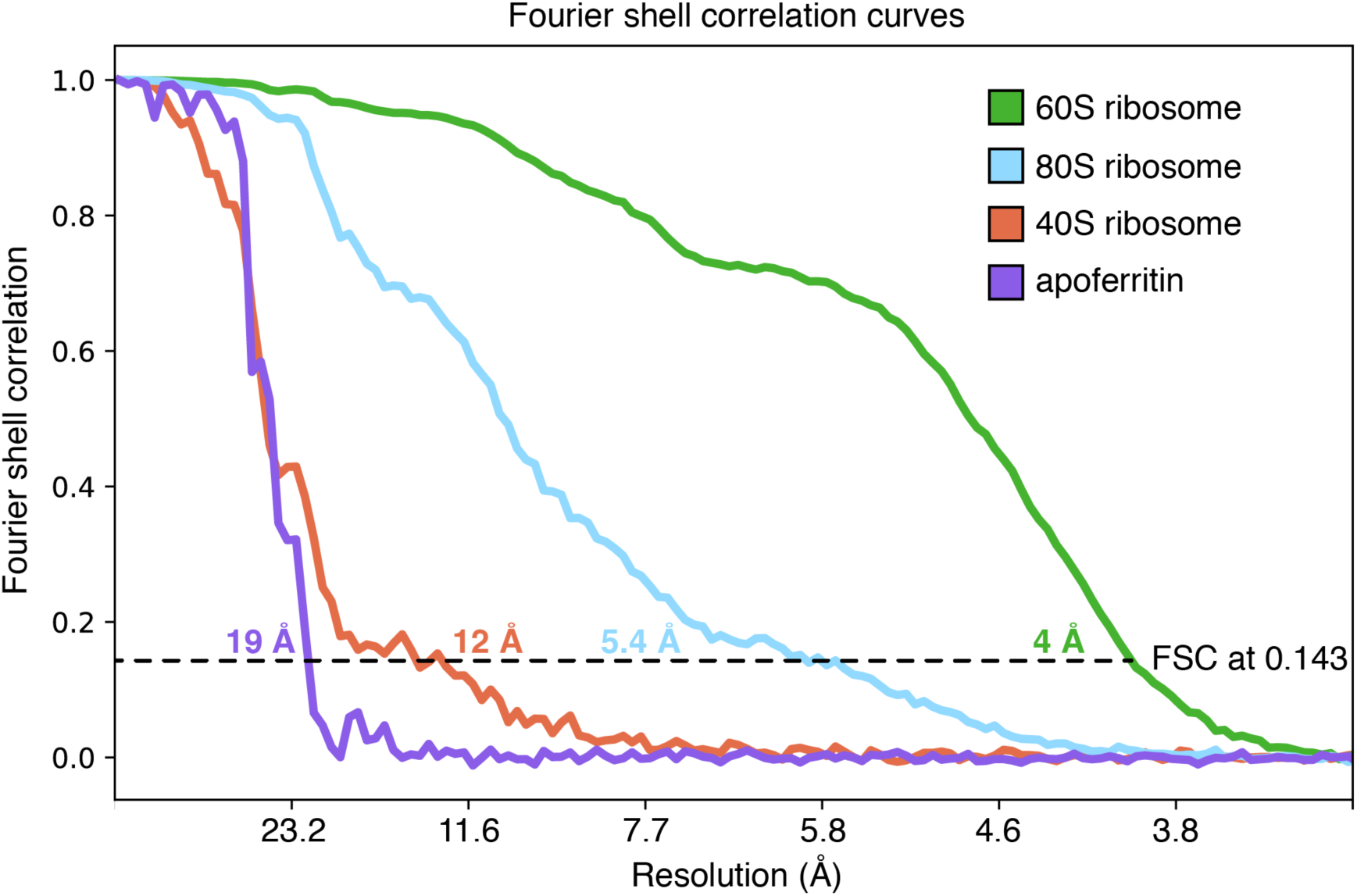
Ab initio reconstructions in cryoSPARC with varying class number. 3D reconstructions using *ab initio* reconstruction in cryoSPARC from the entire data set with K = 3, 4, 5 and 6 classes, respectively.

**Figure S5.**
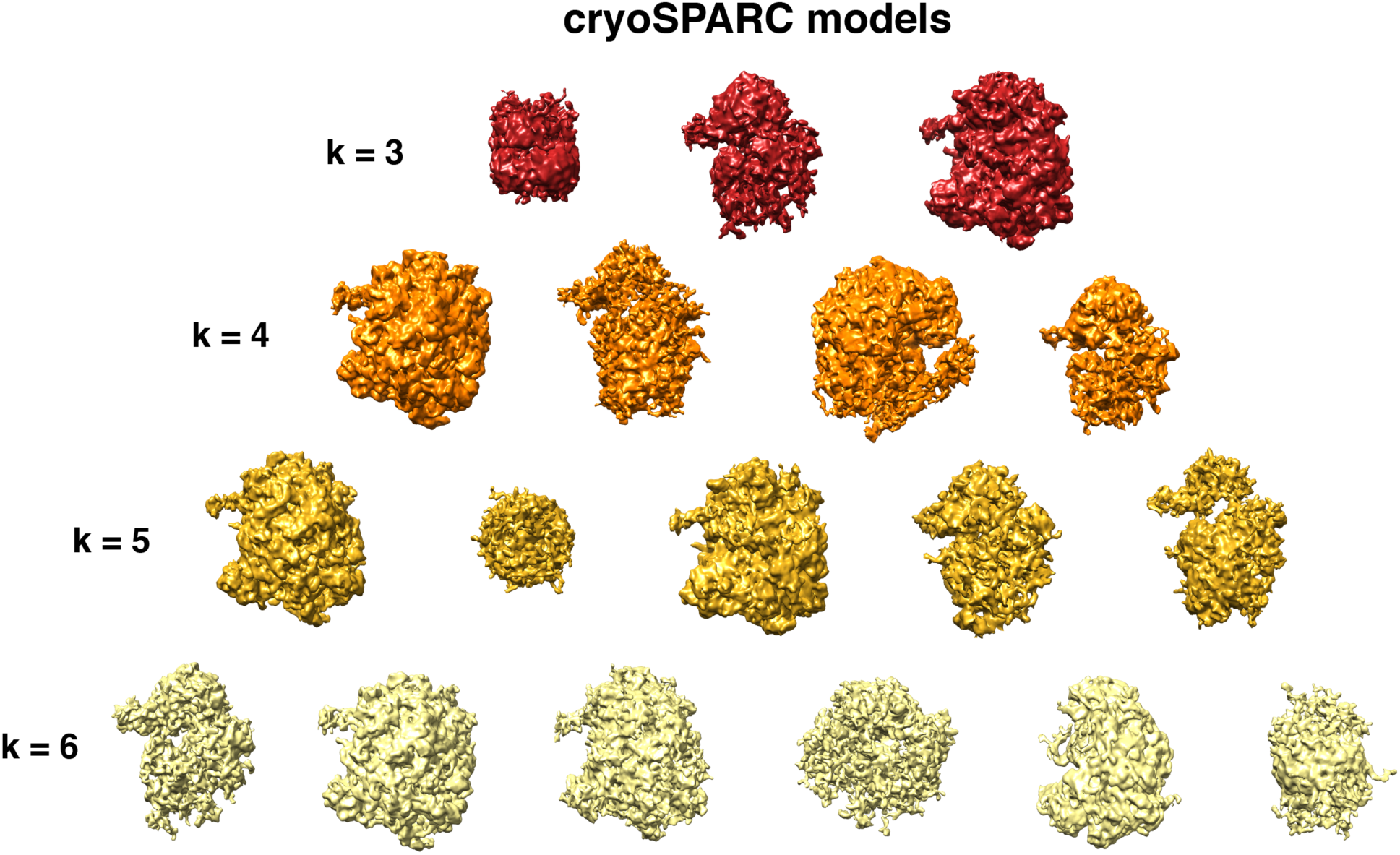
Fourier shell correlations curves. FSC curves for our clustered 80S ribosome (blue), 60S ribosome (green), 40S ribosome (red) and apoferritin (purple) shown in Figure 5B. Nominal resolutions were estimated to be 5.4, 4, 12 and 19 Å, respectively, using the 0.143 gold-standard FSC criterion.

